# ManiNetCluster: A Manifold Learning Approach to Reveal the Functional Linkages Across Multiple Gene Networks

**DOI:** 10.1101/470195

**Authors:** Nam D. Nguyen, Ian K. Blaby, Daifeng Wang

## Abstract

The coordination of genome encoded function is a critical and complex process in biological systems, especially across phenotypes or states (*e.g*., time, disease, organism). Understanding how the complexity of genome-encoded function relates to these states remains a challenge. To address this, we have developed a novel computational method based on manifold learning and comparative analysis, ManiNetCluster, which simultaneously aligns and clusters multiple molecular networks to systematically reveal function links across multiple datasets. Specifically, ManiNetCluster employs manifold learning to match local and non-linear structures among the networks of different states, to identify cross-network linkages. By applying ManiNetCluster to the developmental gene expression datasets across model organisms (*e.g*., worm, fruit fly), we found that our tool significantly better aligns the orthologous genes than existing state-of-the-art methods, indicating the non-linear interactions between evolutionary functions in development. Moreover, we applied ManiNetCluster to a series of transcriptomes measured in the green alga *Chlamydomonas reinhardtii*, to determine the function links between various metabolic processes between the light and dark periods of a diurnally cycling culture. For example, we identify a number of genes putatively regulating processes across each lighting regime, and how comparative analyses between ManiNetCluster and other clustering tools can provide additional insights. ManiNetCluster is available as an R package together with a tutorial at https://github.com/namtk/ManiNetCluster.

## 1. Introduction

The molecular processing between genotype and phenotype is complex and poorly characterized. Understanding these mechanisms is crucial to comprehend how proteins interact with each other in a coordinated fashion to implement various genomic functions and affect high-level phenotypes or biological states (*e.g*., time, disease, organism). Biologically-derived data has undergone a revolution in recent history thanks to the advent of high throughput sequencing technologies, resulting in a deluge of genome and genome-derived (*e.g*. transcriptome) datasets for various phenotypes. Extracting all significant phenomena from these data is fundamental to completely understand how dynamic functional genomics vary between conditions and states such as environment and disease. However, the integration and interpretation of systems-scale (*i.e*. ‘omics’) datasets for understanding how the interactions of genomic functions especially from different gene sets relate to different phenotypes remains a challenge.

Whereas the genome and genes contained are near-static entities within an organism, the transcriptome and proteome are dynamic and state-dependent. The relative quantity of each mRNA and protein species, defining the transcriptome and proteome respectively, function together as networks to implement biological functions. Such networks provide powerful models allowing the analysis of interconnections between biological datasets; *e.g*., gene co-expression networks, derived from transcriptomes, are frequently used to investigate the genotype-phenotype relationships and individual protein function predictions (Carter et al., 2004; Langfelder and Horvath, 2008; Liao et al., 2011; Yang et al., 2014; Zhang and Horvath, 2005). To discover the functional network components, clustering methods have been widely used to detect the network structures that imply functional groupings among genes (*e.g*., gene co-expression modules) (Langfelder and Horvath, 2008). Clustering could be seen as grouping together similar objects; therefore, the key factor to consider first is the distance metric. Previous studies have suggested that some specific distance metrics are only suitable for some certain algorithms and vice versa (Aggarwal et al., 2001; Jaskowiak et al., 2014; Singh et al., 2013; Yang et al., 2016); *e.g*., *k*-means algorithm works effectively with Euclidean distance in low dimensional space but not for high dimensional one such as gene expression datasets (Aggarwal et al., 2001; Yang et al., 2016). More importantly, genes in the network highly likely interact with each other in a non-linear fashion (Yan et al., 2016); many biological pathways involve the genes with short geodesic distances in gene co-expression networks (Yip and Horvath, 2007). To model these non-linear relationships inferring gene function, non-linear metrics including geodesic distance have been used to quantify the similarity between genes and find the non-linear structures of gene networks (*e.g*., manifold) (Lawrence, 2012).

While network analysis is a useful tool to investigate the genotype-phenotype relationships and to derive the biological functional abstraction (*e.g*., gene modules), it is hard to understand the relationships between conditions, and, in particular between different experiments (*e.g*., organisms, environmental perturbations). Therefore, comparative network analyses have been developed to identify the common network motifs/structures preserved across conditions that may yield a high-level functional abstraction. A number of computational methods have been developed to aid biological network, and comparative network analysis (Langfelder and Horvath, 2008; Yan et al., 2014; Zhang and Horvath, 2005). However, these methods typically rely on external information and prior knowledge to link individual networks and find cross-network structures such as counting shared or orthologous genes between cross-species gene co-expression networks (Zeng et al., 2008). Consequently, they potentially miss the unknown functional links that can happen between different gene sets. For example, the genes that express at different stages during cell fate and differentiation can be co-regulated by common master regulators (Lefebvre et al., 2010; Mattick et al., 2010). Additionally, in many cases that the datasets for different conditions are generated independently, individual networks constructed from these datasets of individual potentially have the network structures that are driven by data biases rather than true biological functions. To address this, a comparative method to uniformly analyze cross-condition datasets is essential.

To help overcome some of these limitations, we have developed a manifold learning-based approach, ManiNetCluster, to simultaneously align and cluster multiple gene networks for comparative network analysis. ManiNetCluster enables discovery of inter-network structures implying potential functional linkage across multiple gene networks. This method addresses the challenges for discovering (1) non-linear manifold structures across multiple gene expression datasets and (2) the functional relationships between different gene modules from different datasets. Manifold learning has been successfully used to find aligned, local and non-linear structures among non biological networks; *e.g*., manifold alignment (Ham et al., 2005; Wang and Mahadevan, 2009) and warping (Vu et al., 2012). Previous efforts have resulted in tools that combine manifold learning and gene expression analysis (Narayanan et al., 2010), or to bring together manifold learning and simultaneous clustering (Elhamifar and Vidal, 2011). However, to our knowledge, ManiNetCluster is the first which integrates manifold learning, comparative analysis and simultaneous network clustering together to systematically reveal genomic function linkages across different gene expression datasets. ManiNetCluster is publicly available as an R package at http://github.com/namtk/ManiNetCluster.

ManiNetCluster can be considered as a network embedding method to solve the network alignment problem, which aims to find the structure similarities between two or more (biological) networks. Due to the NP-completeness of the sub-graph isomorphism problem, state-of-the-art network alignment methods often requires heuristic approaches, mapping nodes across networks to maximize a ‘‘topological” cost function, *e.g*., S^3^ (symmetric sub-structure score) measure of static edge conservation (Saraph and Milenkovic, 2014) and static graphlet-based measure of node conservation (Saraph and Milenkovic, 2014; Vijayan et al., 2015), PageRank based cost function and Markovian alignment strategies (Kalecky and Cho, 2018; Liao et al., 2009; Singh et al., 2008). Unlike these topological approaches, which is based on network structure, ManiNetCluster is a (sub)space learning approach, embedding the nodes across multiple networks into a common low dimensional representation such that the distances between mapped nodes as well as the ‘distortion” of each network structure are minimized. ManiNetCluster achieve this goal by implementing manifold alignment (Ham et al., 2005; Wang and Mahadevan, 2009), which is shown in our paper as a manifold co-regularization (Sindhwani and Rosenberg, 2008) method. Because of this manifold regularization (Belkin et al., 2006) nature, our method is a semi-supervised approach since all the mapping of nodes can be propagated from a very few seeds, *i.e*., mappings. Furthermore, the fusion of networks in a common latent manifold allows us to identify not only conserved structure but also functional linkage across networks, a novel type of structure.

## 2. Methods

ManiNetCluster is a novel computational method exploiting manifold learning to discover putative functional linkages across multiple gene networks (Figure 1, Algorithm 1). By importing multiple gene expression datasets across conditions such as phenotypes or states, the tool constructs the gene networks for each state, in which genes are connected if the similarity of their expression profiles for the state is high (*i.e*., co-expression). The multiple gene networks can be interconnected using the same genes (if the datasets are across conditions) or orthologs (if the comparison is between two organisms). Secondly, ManiNetCluster uses manifold alignment (Ham et al., 2005; Wang and Mahadevan, 2009) or warping (Vu et al., 2012) to align multiple gene networks (*i.e*., to match their manifold structures that are typically local and non-linear across time points), and assemble these aligned networks into a multilayer network. In particular, this alignment step projects two high-dimensional gene networks into a common lower dimensional space on which the Euclidean distances between genes preserve the geodesic distances that have been used as a metric to detect manifolds embedded in the original high-dimensional ambient space (Belkin and Niyogi, 2003). Finally, ManiNetCluster simultaneously clusters this multilayer network into a number of cross-network gene modules. These gene modules can be characterized into: (1) the conserved modules mainly consisting of the same or orthologous genes; (2) the condition-specific modules mainly containing the genes connected from one network; (3) the cross-network linked modules consisting of different gene sets from each network and limited shared/orthologous genes (Figure 1). In particular, we refer to the latter module type as the ‘functional linkage” module. This module type demonstrates that different gene sets across two different conditions can be still clustered together by ManiNetCluster, suggesting that the cross-condition functions can be linked by a limited number of shared genes. Consequently, and more specifically, these shared genes are putatively involved in two functions in different conditions. These functional linkage modules thus provide potential novel insights on how various molecular functions interact across conditions such as different time stages during development.

**Figure 1:**
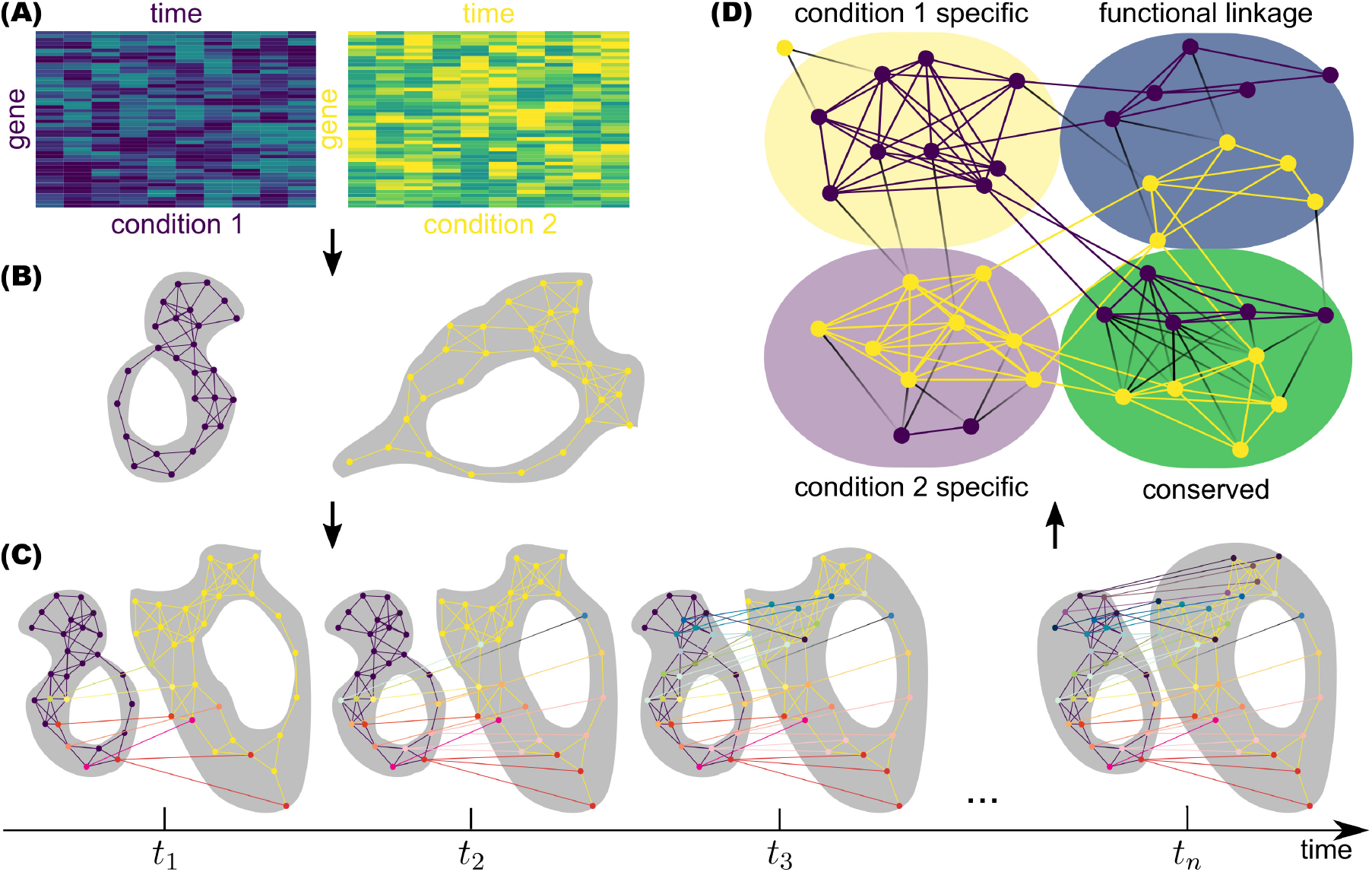
ManiNetCluster Workflow Schematic. (A) The inputs of the workflow are the two time series gene expression datasets collected from different conditions (as in this example) or from two different organisms. The number of genes and/or the number of timepoints need not be the same in each dataset. (B) From the input data, gene co-expression neighborhood networks are constructed, which approximate the manifolds where the datasets are concentrated. (C) Using manifold alignment and manifold warping methods, the two gene expression profiles are aligned across time series in a common manifold. The outcome of this step is a multilayer network consisting of two types of links: the inter-links showing the correspondence (*e.g*. shared genes) between the two datasets, and the intra-links showing the co-expression relationships. D) The multilayer network is clustered into modules. Four distinct types of modules are revealed in this step: conserved modules containing high proportion of shared genes, dataset 1-specific modules containing a high proportion of genes from dataset 1, dataset 2-specific modules containing a high proportion of genes from dataset 2, and function linkage modules containing near equal numbers of genes from both datasets, which are the same gene (if conditions are compared from the same organism) or orthologs (if the organism differs between the two compared datasets).

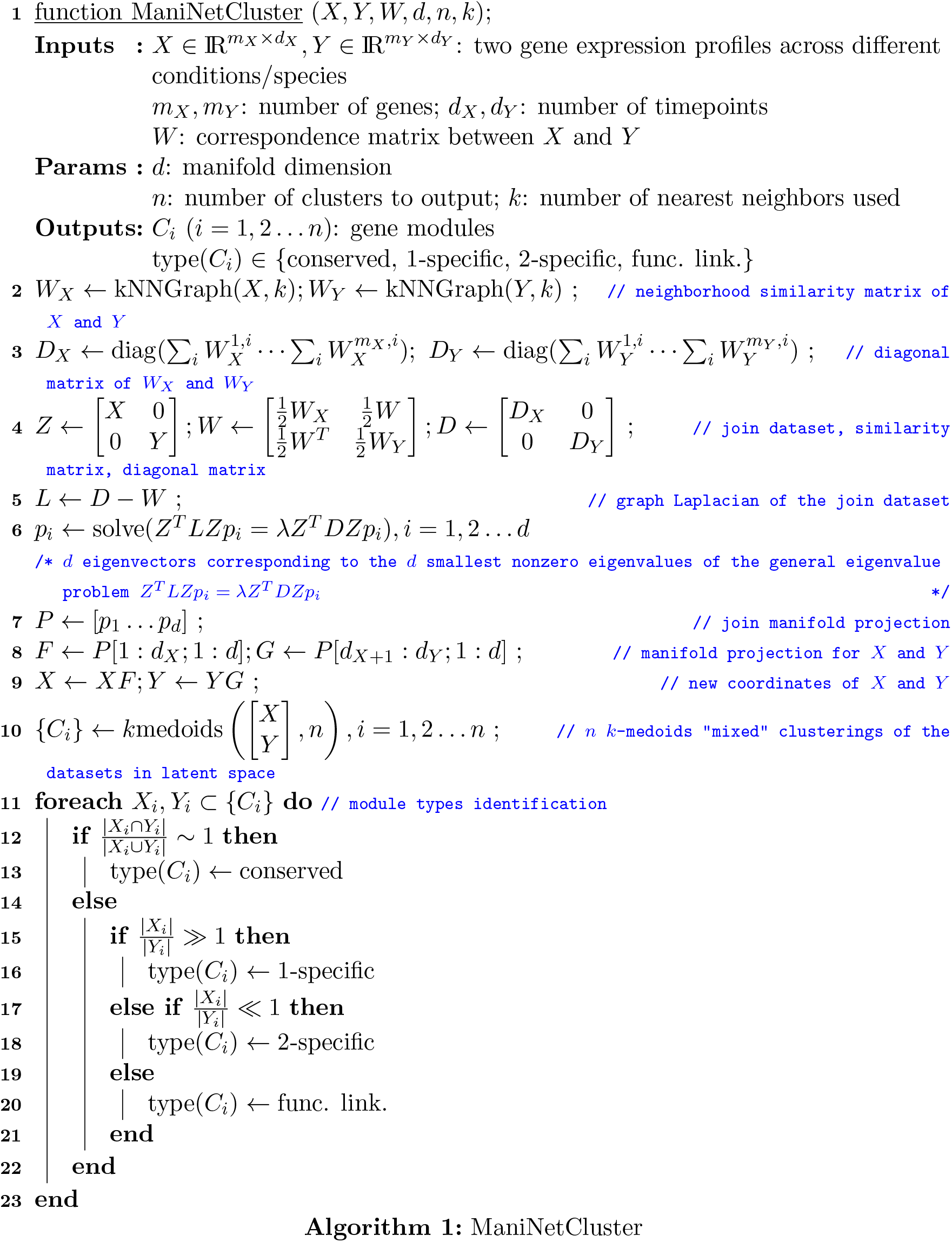

A detailed overview of ManiNetCluster is depicted in Algorithm 1. Step 1 is problem formulation. The next steps describe the primary method, which can be divided into two main parts: steps 2 to 9 are for manifold alignment; steps 10 to 20 are for the simultaneous clustering and module type identification. Our method is as follows: first, we project the two networks into a common manifold which preserves the local similarity within each network, and which minimizes the distance between two different networks. Then, we cluster those networks simultaneously based on the distances in the common manifold. Although there are some approaches that use manifold alignment in biological data (Alpert et al., 2018; Welch et al., 2017), our approach is unique since it deals with time series data (when using manifold warping) and the criteria that lead to the discovery of four different types of functional modules. The details of the two main parts are as follows.

### 2.1 Manifold alignment/warping

The first steps of our method (steps 2 to 9) are based on manifold alignment (Wang and Mahadevan, 2009) and manifold warping (Vu et al., 2012), which based on the manifold hypothesis, describes how the original high-dimensional dataset actually lies on a lower dimensional manifold, which is embedded in the original high-dimensional space (Fefferman et al., 2016). In ManiNetCluster, we project the two networks into a common manifold which preserves the local similarity within each network and which minimizes the distance between two different networks.

We take the view of manifold alignment (Wang and Mahadevan, 2009) as a multi-view representation learning (Wang et al., 2015), in which the two related datasets are represented in a common latent space to show the correspondence between the two and to serve as an intermediate step for further analysis, *e.g*. clustering. In Algorithm 1, we described the parametric manifold alignment where the transformation *F* and *G* can be learned explicitly from two disparate gene expression profiles. In general, given two disparate gene expression profiles 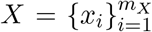 and 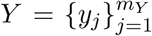 where *x_i_* ∈ ℝ^*d_X_*^ and *y_j_* ∈ ℝ^*d_Y_*^ are genes, and the partial correspondences between genes in *X* and *Y*, encoded in matrix *W* ∈ ℝ^*m_X_ × m_Y_*^, we want to learn the two mappings *f* and *g* that maps *x_i_, y_j_* to *f*(*x_i_*), *g*(*y_j_*) ∈ ℝ^*d*^ respectively in a latent manifold with dimension *d* ≪ *min*(*d_X_, d_Y_*) which preserves local geometry of *X, Y* and which matches genes in correspondence. We then apply the framework in vector-valued reproducing kernel Hilbert spaces (Minh et al., 2016; Minh and Sindhwani, 2011) and reformulate the problem as follows to show that manifold alignment can also be interpreted as manifold co-regularization (Belkin et al., 2006).

Let *f* = [*f*_1_… *f_d_*] and *g* = [*g*_1_…*g_d_*] be components of the two ℝ^*d*^-value function *f*: ℝ^*d_X_*^ → ℝ^*d*^ and *g*: ℝ^*d_Y_*^ → ℝ^*d*^ respectively. We define Δ*f* ≜ [*L_X_ f*_1_ … *L_X_ f_d_*] and Δ*g* ≜ [*L*_*Y g*_1__…*L_Y g_d__*] where *L_X_* and *L_Y_* are the scalar graph Laplacians of size *m_X_* × *m_X_* and *m_Y_* × *m_Y_* respectively. For 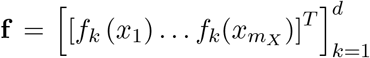 and 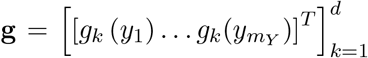, we have 〈**f**, Δ_*X*_**f**)_ℝ^*dm_X_*^_ = *trace*(**f**^*T*^*L_X_***f**) and 〈**g**, Δ_*Y*_**g**〉_ℝ^*dm_Y_*^_ = *trace*(**g**^*T*^*L_Y_***g**). Then, the formulation for manifold alignment is to solve,

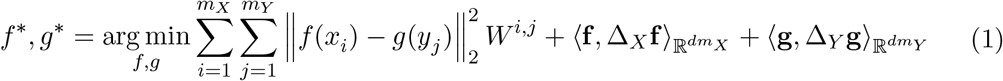

The first term of the equation is for obtaining the similarity between corresponding genes across dataset; the second and third terms are regularizers preserving the smoothness (or the local similarity) of the two manifolds. In parametric approach, finding minimizers *f*\* and *g*\* is equivalent to finding the solution of the general eigenvalue problem *Z^T^LZ_pi_* = λ*Z^T^DZ_pi_* where 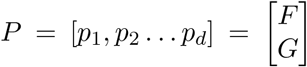 and *XF* = **f**, *YG* = **g** as depicted in steps 6 to 9 of Algorithm 1. Manifold alignment can also be non-parametric where, instead of finding linear form of transformation *F* and *G* from steps 6 to 9, we find the new coordinates *X*’ and *Y*’ directly by solving the general eigenvalue problem *L_pi_* = λ*D_pi_* where 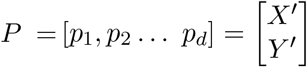 and *X*’ = **f**, *Y*’ = **g**

In biological settings, the two disparate datasets *X, Y* share the similar underlying manifold representation because they are gene expressions from different conditions yet of the same species, or in other case, from different species yet of the same branch of evolutionary tree. From these two gene expression profiles, two gene co-expression neighborhood networks are implicitly constructed as approximations of the two manifolds. Then, the two manifolds are aligned providing the pairwise correspondence between the two datasets *W* according to the optimization problem in equation 1. The correspondence matrix *W* could be an identity matrix if the problem is cross-condition analysis within a specific species or could be the one whose elements 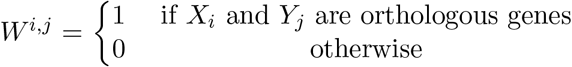 if the problem is cross-species analysis. Alternatively, in manifold warping (Vu et al., 2012), the correspondence matrix *W* is not provided but learned with time warping function. As a result, this gives us two transformed datasets where the pairwise distance among the two dataset is diminished (compared to the original dataset).

### 2.2. Simultaneous clustering and characterization of module types

Our ultimate goal is to simultaneously cluster the genes across different conditions so that we can actively detect which modules are conserved, which modules are specific and most importantly, which modules are functional linkage. To obtain such results, we deal with two challenges, which are (1) to integrate data across different conditions in a meaningful way and (2) to come up with a suitable distance measurement. Using manifold alignment/warping methods, we could solve those two problems together, since in manifold alignment the two datasets are projected into the latent common space where distances between corresponding points are minimized and where the locality could be measured using Euclidean distance. Thus, we perform the clustering on top of the transformed data, in which the transformation is calculated in the previous step using manifold alignment/warping methods. We applied *k*-medoids clustering for the robustness over outliers and obtained the modules whose genes might be of either of the two original networks; the proportion of such genes between networks inside a module would tell the type of that module: conserved, condition 1-specific, condition 2-specific, or functional linkage.

Simultaneously clustering is performed over the concatenation of transformed datasets: Two disparate datasets are embedded in a common latent manifold, whose geodesic distances between points are preserved. The concatenation of the embedded datasets 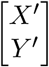 where *X*’ = *XF, Y*’ = *YG* are then simultaneously clustered (using *k*-medoids). The clustering is shown in step 10 of the Algorithm 1.

We then identified two criteria to delineate the four types of genomic functional modules, which are conserved modules, data 1 specific modules, data 2 specific modules, and functional linkage modules: (1) the so-called Condition number, which is the fraction between number of genes from dataset 1 over the number of genes from dataset 2, and (2) the so-called intra-module Jaccard similarity between the the two gene sets from two conditions. Concretely, the clustering results *C*_1_, *C*_2_ …*C_n_* (gene modules) are of 4 types, characterized by intra-module Jaccard similarity 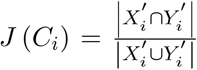 and Condition number 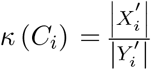. The criteria are depicted in Algorithm 1 from step 11 to step 20

In our method, the cross-network functional linkage module is the conjugate of the conserved module in this following way: while the intra-module Jaccard similarity in conserved modules is high because the corresponding genes in both datasets appear in the same cluster, the intra-module Jaccard similarity in functional linkage module is low. In short, we use Euclidean distance embedded locally in the manifold for clustering and intra-module Jaccard similarity for detecting type of modules. This type of modules consisting of different genes from multiple networks/conditions that are clustered together from ManiNetCluster, revealing potential cross-condition functional linkages between different gene sets, *e.g*. photosynthesis function linkages.

We argue the existence of functional linkage modules as follows. The two datasets are transformed and lay in a latent common space where the Euclidean distance could be used locally to illustrate the similarity between close points (genes) and the Jaccard similarity is used to encode the similarity of the two datasets in terms of correspondence information. In the case of functional linkage module, the genes are close but not corresponding to each other, implying that the genes are similar in functions but still different since they belong to different parts of the two datasets.

## 3. Experiments and results

### 3.1 Datasets

To validate our methods, we applied ManiNetCluster to several previously published datasets:

1. *Developmental gene expression datasets for worm and fly:* The dataset describes time-series gene expression profiles of *Caenorhabditis elegans* (worm) and *Drosophila, melanogaster* (fly), taken during embryogenesis developmental stage. The data is from the comparative modENCODE Functional Genomics Resource (Celniker et al., 2009). We took 20377 genes over 25 stages for worm and 13623 genes over 12 timepoints for fly. After removing low expressed genes, we were left with 18555 and 11265 genes for worm and fly respectively. From these genes, we took 1882 fly genes and 1925 worm genes which have orthologous as correspondence information for our alignment methods (Celniker et al., 2009). The gene expression data is then normalized to unit norm.
2. *Time-series gene expression datasets for alga:* This dataset describes the transcriptome in a synchronized microalgal culture grown over a 24hr period (Zones et al., 2015). The data contains 17737 genes over 13 timepoints sampled during the light period and 15 timepoints sampled during the dark period. After removing low expressed genes, we were left with 17695 for further analysis with ManiNetCluster.

To remove noise from gene expression data signal, we removed genes whose expression value was less than 1 across all time points. Also, we looked for any outliers in the datasets by hierarchical clustering across all time points. This resulted in omitting 42 low expressed genes as for the alga dataset, 1822 worm genes and 2358 fly for worm and fly datasets; there were no outliers in any of the datasets used. Next, log2 transformed of the algal dataset.

### 3.2. ManiNetCluster reveals conserved manifold structures between cross-species gene networks

In addition to being able to cluster co-expressed genes, a unique aspect of ManiNetCluster is the ability to directly identify which modules are conserved, specific, putatively functionally linked without further analysis. ManiNetCluster organizes genes into clustered modules using a manifold alignment/warping approach. Unlike other hierarchical or *k*-means methods for clustering, our platform enables the simultaneous clustering of different datasets, offering the possibility of novel biological insight via the comparison of multiple independent experiments. This is due to the simultaneous clustering of datasets, whereas other clustering methods treat each gene expression dataset derived under different conditions separately. This uniquely allows for the identification of groups of genes, potentially linked biologically, that would otherwise be missed, possibly elucidating novel phenomena or functional inferences.

We previously demonstrated that orthologs across multiple species function similarly in development by using a networking approach (Celniker et al., 2009; Yan et al., 2014).

However, not all orthologs have correlated developmental gene expression profiles (Singh et al., 2008), suggesting that they may have non-linear relationships in terms of gene expression. To investigate this discrepancy, we applied ManiNetCluster to the time-series gene expression datasets of model organisms, *Caenorhabditis elegans* (worm) and *Drosophila melanogaster* (fly), taken during embryogenesis, to determine whether orthologous genes have non-linear relationships, and if these relationships are also conserved across species. We employed ManiNetCluster to align cross-species developmental gene networks. These analyses indicated that the orthologous genes between worm and fly are better aligned by non-linear manifold learning than the linear methods, as indicated by their distances after alignment: CCA = 632.44 *vs*. ManiNetCluster = 276.32 (t-test p-value < 2.2 × 10^−16^) in terms of sum of pairwise distances (*i.e*., less distance and more preserved network structure after alignment; Figure 2). This suggests that non-linear interactions exist between evolutionary conserved functions encoded by orthologues genes across worm and fly during development.

**Figure 2:**
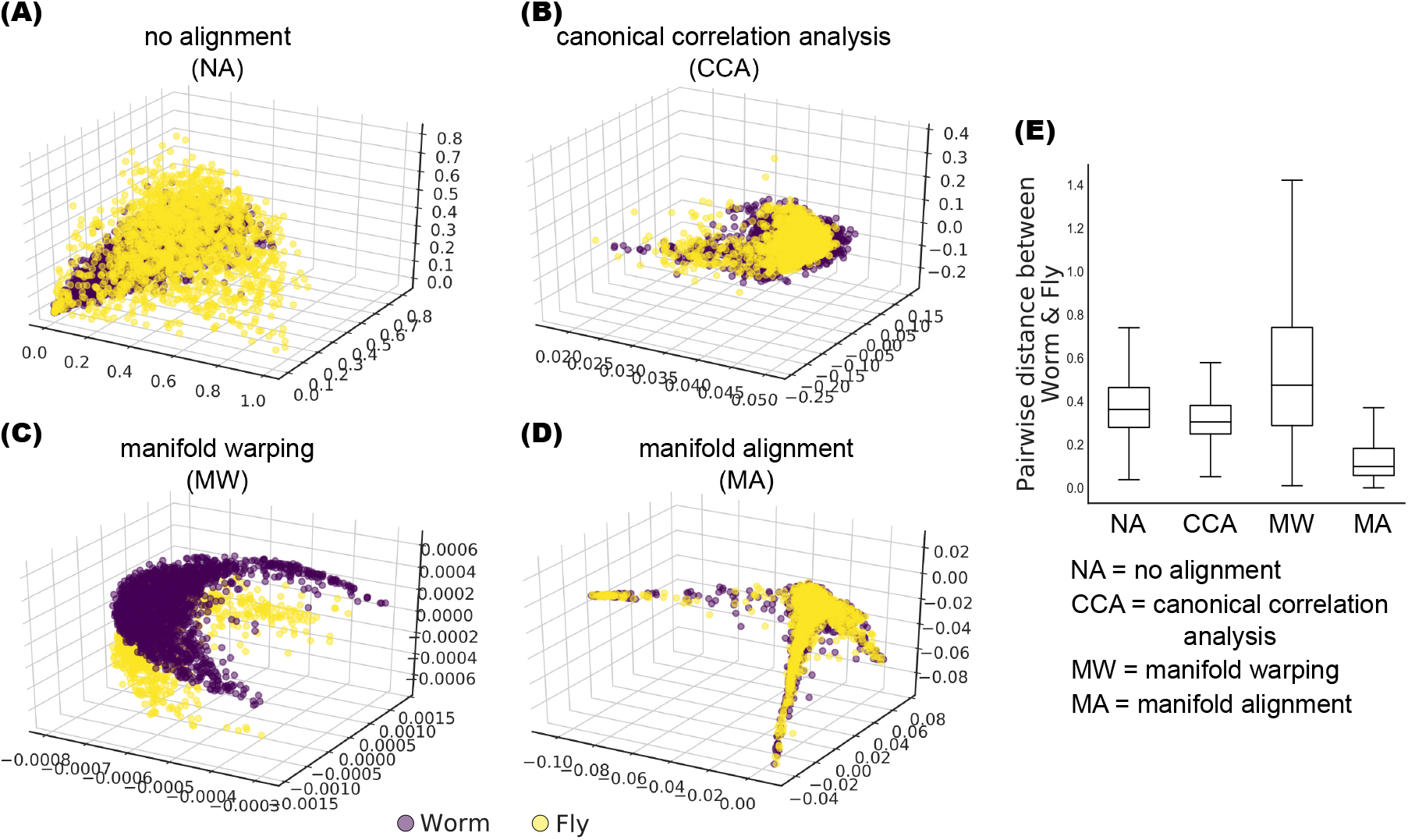
ManiNetCluster outperforms alternative methods to align and identify non-linear structures between cross-species developmental gene networks. Absence of data alignment, canonical correlation analysis, manifold warping and manifold alignment methods of alignment are shown using worm (purple) and fly (yellow) datasets in A-D respectively. (E) Boxplots depicting the gene distance between two species (using Chebyshev distance) in all alignment methods. The box extends from the lower to upper quartile values of the data (pairwise distance between worm and fly), with a line at the median. The whiskers extend from the box to show the range of the data. Outliers beyond the whiskers are omitted from the plot.

### 3.3 ManiNetCluster identifies putative genomic function links between cross-condition gene networks

As a case study to demonstrate the uniqueness and validity of ManiNetCluster for comparing between conditions, we used a previously published dataset (Zones et al., 2015). This dataset describes the transcriptomic dynamics of a synchronized microalgal culture grown over a 24hr period, and was specifically chosen to test ManiNetCluster due to the comprehensiveness of the time series (samples taken at 1 hour or 30 minute intervals over two independent 24 hour periods (Zones et al., 2015)). Using the ManiNetCluster algorithm we delineated the transcriptomes sampled during the light period vs. the dark period of the 24 hour experiment. After alignment (in which ManiNetCluster again outperformed CCA: ManiNetCluster = 128.00 *vs*. CCA = 713.50 in terms of sum of pairwise distances (t-test p-value < 2.2 × 10^−16^)), we simultaneously clustered the two groups of transcriptomes, treating the light- and dark-collected samples as independent experiments. ManiNetCluster clustered the two datasets (*i.e*. light period and dark period) into 60 modules of *Chlamydomonas reinhardtii*, and delineated the genes in each into light-specific, dark-specific and shared between light and dark (Figure 3; Supplemental Tables 1 & 2). Based on the metrics (intra-module Jaccard similarity, Condition number) that quantify relative light/dark gene proportions (Methods; Supplemental Table 2), we detected four types of module: conserved, light or dark specific, and functionally linked. The functional linkage modules consist of different gene sets from light and dark networks with very limited shared genes (Supplemental Table 2). For example, Module 60 is a dark-specific module due to a high proportion of dark period genes and Module 21 is a conserved module since it has a high fraction of shared genes (Figure 3; Supplemental Tables 1 & 2). Module 34 is a functional linkage module since it contains a low proportion of shared genes and high proportion of different light and dark period genes (Figure 3; Supplemental Tables 1 & 2). Many modules are highly enriched for genes expressed during the light period, the dark period and for shared in both the light and dark networks. This is clearly demonstrated in Modules 34, 52 and 60, which are enriched for shared, light and dark genes respectively (Figures 3 & 4; Supplemental Tables 1 & 2). These groupings indicate that the proteins encoded by genes in these modules could have related specific roles in either light-, dark-or both light and dark-specific metabolism. Consequently, the gene sets within each module could be used to provide functional inferences for each gene and the co-expressed genes across the module. For example, Module 21 is highly enriched for genes encoding proteins involved in protein synthesis in the light-dark shared fraction of the module, suggesting that these proteins are active in the synthesis of proteins for both the light and dark periods.

**Figure 3:**
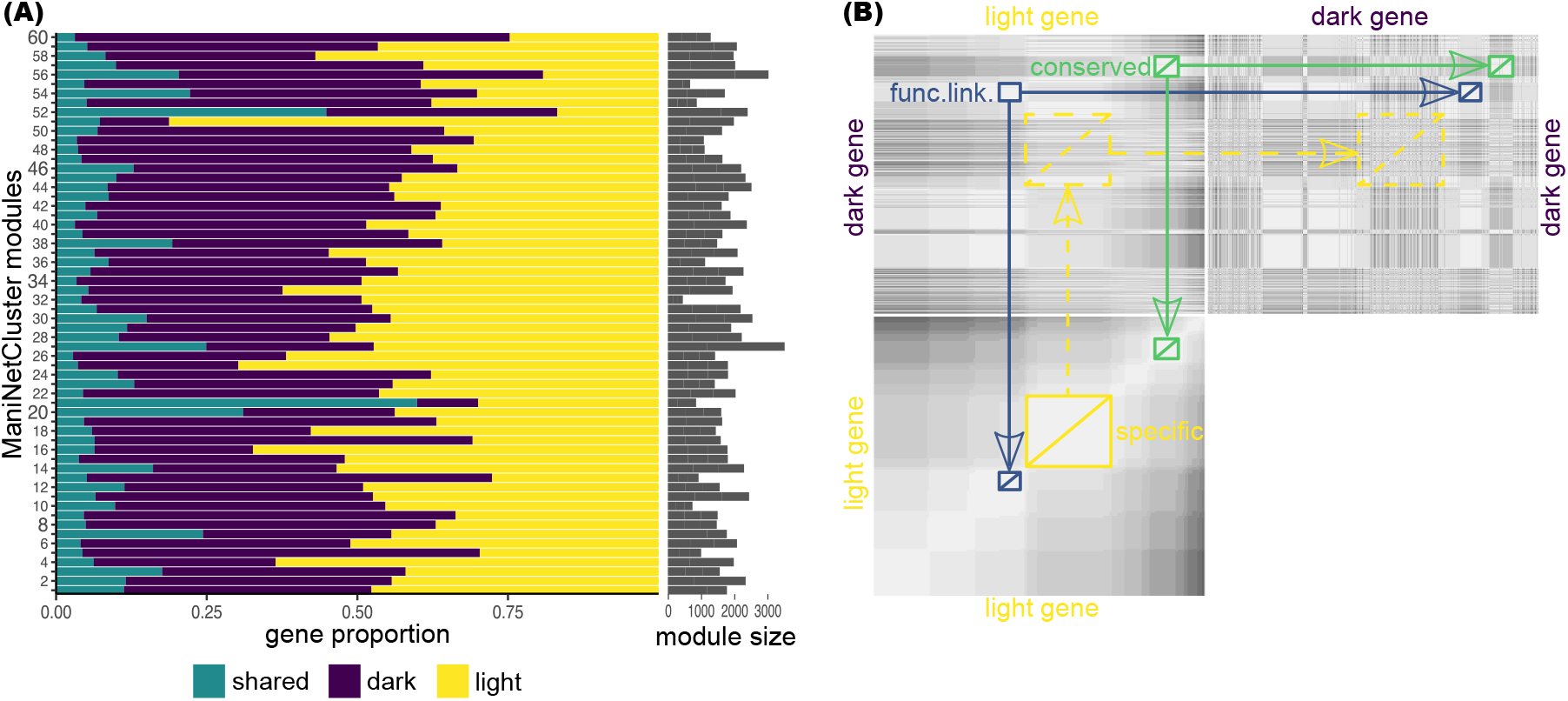
Clustering Results of ManiNetCluster, and identification of different module types. (A) Clustering of algal time series diurnal transcriptomes \cite{RN21} into 60 modules using ManiNetCluster. For the purposes of these analyses, the transcriptomes collected during the light period were treated as an independent experiment from those collected during the dark period. The proportion of each module comprised of light period specific (yellow) dark period specific (purple) and shared (teal) is shown. Module size is indicated on the right of the modules. Complete module data are shown in Supplemental Tables 1 and 2. (B) Cross-heatmap demonstrating the relationship between modules in each condition (*i.e*. light period-specific or dark period-specific), which reveals the module types. The off-diagonal module (depicted in blue) which has corresponding modules in both light and dark clusters is an example of a functionally linked module, and the on-diagonal module (depicted in green) which has corresponding modules in both light and dark clusters is an example of condition-specific module.

To further investigate and validate the functional linkage modules, we focus here specifically on two Modules, 6 and 34 (Figures 3 and 4; Supplemental Tables 1 & 2). These modules were chosen as examples since they both exhibit low intra-module Jaccard similarities (0.04 and 0.03 for Modules 6 and 34 respectively) and their Condition number values is approximately 1 (1.13 and 1.04 for Modules 6 and 34 respectively), indicative of a small number of shared genes and similar numbers of light and dark period genes (Supplemental Table 2). Module 34 contains a total of 598 genes. Of these, the mRNA abundance of 284 genes within the module are from the light period and 295 are from the dark period (Figures 3 and 4; Supplemental Table 1). Of those genes annotated, the light period genes are functionally enriched for flagellar associated proteins (FAPs (Pazour et al., 2005)), the cell motility and cell organization Mapman ontologies (Thimm et al., 2004) and the dark period genes contain a number of transporters, Greencut associated genes (Heinnickel and Grossman, 2013; Karpowicz et al., 2011; Merchant et al., 2007) and genes encoding proteins involved in DNA synthesis. More notably, 19 genes are shared between the light and dark periods, meaning that these genes tightly co-express with both the light genes during the light period and the dark genes during the dark period (Figure 4; Supplemental Table 1). These 19 genes encode proteins functionally enriched for aspects of regulation, including protein post-translational modification and RNA regulation (8 of the 19 genes have an associated gene ontology, all of which are related to regulation. These ontologies (and gene annotations where they exist), together with the interactions with the rest of the module, suggest the possibility of a hierarchical gene/protein regulatory network, with these genes putatively imposing some aspect of regulation upon the rest of the module. Similarly, Module 6 contains 721 genes, of which 326 are dark-period specific, 368 are light-period specific and 27 are shared. Again, these 27 are enriched for genes encoding proteins with putative regulatory roles (Figure 4; Supplemental Table 1). Additional modules that display the same statistical characteristics are Modules 15 and 40 (as indicated by the intra-module Jaccard similarities and Condition numbers; Figure 4, Supplemental Table 2).

**Figure 4:**
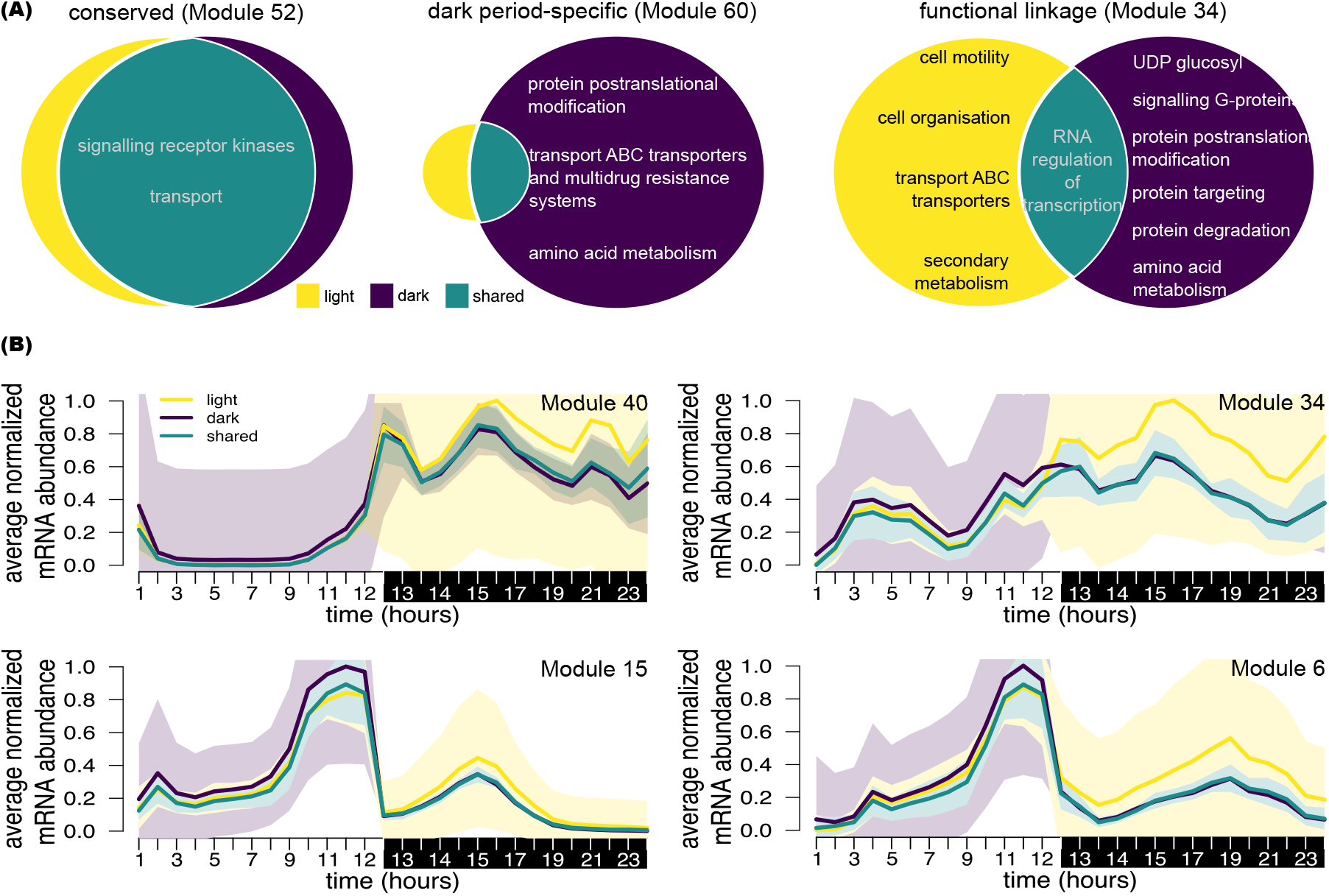
ManiNetCluster identifies different module types. (A) Module types identified by ManiNetCluster, using an algal diurnal dataset (Zones et al., 2015) with light-period and dark-period transcriptomes treated as independent experiments. Example modules are shown: (1) Module 52 - a conserved module in which the proportion of shared genes is high (2) Module 60 - a dark specific module in which the proportion of dark period genes is high, (3) Module 34 - a functional linkage module in which the proportion of shared genes is low and the proportion of light period genes and dark period genes are approximately equal. Functional enrichment for each were generated using MapMan ontologies (Thimm et al., 2004). Complete module data are shown in Supplemental Tables 1 and 2. (B) Expression patterns of example functionally linked modules: Expression patterns of light, dark, and shared genes of modules 34, 6, 40 and 15 are shown. The shared genes (shown in teal) correlate with light genes (purple) in light (13 first time points) and with dark genes (yellow) in dark (15 last time points). The light and dark periods are shown with shading on the x axis. Complete module data are in Supplemental Tables 1 and 2.

## 4. Evaluation

### 4.1 Biological validation

As for validation, we also looked at the biological functions and pathways in ManiNetCluster modules to see if they are consistent with the previous experimental findings. In particular, we tested the biological validity of the modules generated by ManiNetCluster by comparing to modules generated by *k*-means on the diurnal datasets published by Zones et al due to the dataset comprehensiveness outlined above (Zones et al., 2015) (Section 3.1 and 3.2). In that study, using the *k*-means algorithm, 12,592 genes were clustered into co-expressed modules. Since this number represents > 70% of the genes on this organisms’ genome, we reasoned such a significant number would provide an appropriate testbed for corroborating our method described here. The two methods of module generation performed on the same original dataset are highly similar, indicating the general validity of the ManiNetCluster approach in terms of biological significance. Firstly, there is a high degree of similarity of coclustered genes between modules generated using ManiNetCluster and the *k*-means method (ARI = 0.95 and 0.95 for light and dark period modules respectively, see Section 4.2). Secondly, genes encoding proteins of related function are co-expressed, since interacting proteins are required together and under the same conditions.

Analysis of the modules generated by ManiNetCluster indicates functionally-related genes are co-clustered, as expected. For example, the genes encoding proteins constituting the photosynthetic complexes LHCI, LHCII, PSI, PSII, *b*_6_f and the chloroplast ATP synthase are nearly entirely contained within the ManiNetCluster Modules 20 and 21 (Supplemental Table 1). Equally, the genes encoding subunits of the mitochondrial respiratory complexes are almost entirely contained within two modules (Supplemental Table 1), as are the genes encoding many other functionally-related proteins (Supplemental Table 1). Together, these two analyses serve to confirm the veracity of our method for clustering similarly expressed genes.

### 4.2 Comparison of ManiNetCluster *vs.* other clustering methods

Finally, we compared ManiNetCluster to the state-of-the-art methods, including WGCNA, *k*-means, Hierarchical Clustering (HC), Expectation Maximization (EM) that cluster individual gene networks into modules to evaluate the consistency of our clustering. (The technical details of these other methods are specified in Appendix A.1.) As a measure of evaluation, we employed the adjusted rand index (ARI) to assess the overlap of gene modules from these other methods (Figure 5). Specifically, the similarity between two data clusterings *C* = {*C*_1_, *C*_2_…*C_k_*} and 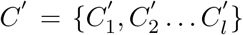 is computed using the adjusted rand index (ARI) as follows:

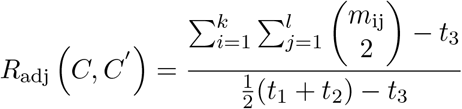

where 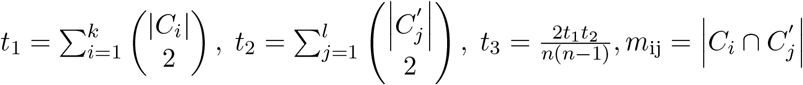, and *n* is the number of observations (*i.e*. genes). The value of this index is ranged from 0 (independant clusterings) to 1 (identical clustering). For this assessment, we again used the datasets from a previously published time series RNA-seq experiment (Zones et al., 2015). Using this data, we found that in general, the ManiNetCluster modules overlap with those identified by other methods (*e.g*., WGCNA = 0.92 and 0.93, *k*-means = 0.95 and 0.95, EM = 0.81 and 0.79, HC = 0.70 and 0.78 for light and dark modules, respectively). The high value of ARI over *k*-means and WGCNA indicates that ManiNetCluster is effective (consistent to *k*-means clustering, proved to deliver meaningful biological results in previous experiment (Zones et al., 2015)) and robust (consistent to WGCNA). This demonstrates that the crosscondition modules of ManiNetCluster are highly consistent with those found by state-of-art methods but more importantly, they provide additional insights into the connections among various genomic functions across different conditions.

**Figure 5:**
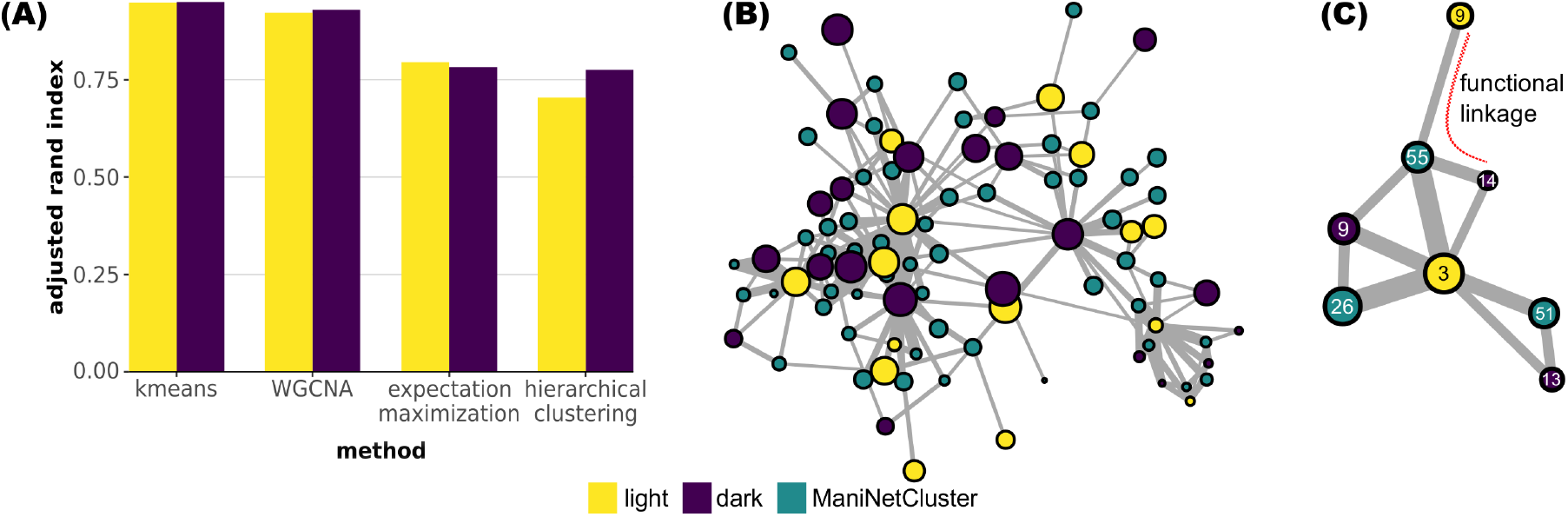
Comparison of ManiNetCluster with other methods. (A) The adjusted rand index between ManiNetCluter clustering and other methods, as shown, indicates ManiNetCluster is consistent with *k*-means and WGCNA but less so with expectation maximization and hierarchical clustering. (B) comparison of 60 cross-condition modules detected by ManiNetCluster and 34 light period modules and 30 dark period modules separately detected by WGCNA by constructing a network, consisting all ManiNetCluster and WGCNA modules as nodes. The links between two nodes indicate the genes shared by both modules. Node size indicates the degree of that node. Links with very low weight are omitted. The triad of the network among three different kinds of nodes (*i.e*. ManiNetCluster module, WGCNA “light-period” module and WGCNA “dark-period” module) indicates the functional linkage type of an ManiNetCluster module. An open triad patterns indicates a functional linkage module. (C) Subgraph of the network in (B) demonstrating a functional linkage module (Module 55). The subgraph also identifies a putative functional link between two WGCNA modules, Light-Module 9 and Dark-Module 14.

To demonstrate this capability, we compared the ManiNetCluster clustering results with those collected using WGCNA to evaluate how they overlap, potentially providing additional functional linkages via this comparative analysis. Specifically, we connected the modules of WGCNA and ManiNetCluster where they share genes, and created a module network in which edge weights are the number of shared genes (Figures 5B and 5C). We found that functional linkage modules generated by ManiNetCluster can connect multiple WGCNA modules (Figure 5). We thus investigated the triad patterns (among ManiNetCluster modules, WGCNA modules for light, WGCNA modules for dark) of such network to analyze if a ManiNetCluster module is of functional linkage type, which is correspondent to the opened triangle (depicted by opened red curve) shown in Figure 5C. For example, Module 55 contains a total of 233 genes, of which 10 are co-expressed with both the light and dark period genes across the complete 24 hour experiment (Supplemental Table 1). Within the 10 shared genes are FTSY, which has a demonstrated role in LHC assembly (Kirst et al., 2012) suggests the possibility of additional roles during the dark period. Another gene in this group is FDX7, encoding a predicted uncharacterized ferrodoxin (Sawyer and Winkler, 2017), suggestive of a role in both the light and dark periods for this protein also. The triad pattern shown in Figure 5C also suggests a functional link between WGCNA Light-Module 9 and WGCNA Dark-Module 14, which cannot be detected by WGCNA itself, since they have shared genes with a ManiNetCluster functional linkage module (Module 55).

## 5. Discussion

Elucidating and understanding the data encoded within each organism’s genome remains the greatest challenge in modern biology. To help extract more information from gene expression datasets, we have developed a novel computational method, ManiNetCluster, which aims to reveal functional linkages of multiple gene networks across conditions (*e.g*., species, time points). In particular, this method extends the manifold learning approaches that capture non-linear relationships among genes to simultaneously cluster multiple gene networks to discover cross-network gene modules linking various genomic functions together. For instance, our tool could be used interrogate two transcriptomes investigating the gene expression effects of two different drug treatments, possibly aiding in the identification of synergistic or antagonistic consequences of dual delivery.

As a tool, ManiNetCluster falls within an emerging field of research, called multi-view learning (Sun, 2013; Xu et al., 2013). Many biological datasets are naturally comprised of different representations or views, which often provide compatible and complementary information (Li et al., 2016a), *e.g*., light and dark period transcriptome of an alga, gene expression of worm and fly whose genes are orthologous or multi-omics single cell data (Colom-Tatch and Theis, 2018). It is natural to integrate these views together (in a nonlinear way) prior to any analysis rather than analyzing each view separately, and then concatenating them (in a linear way). ManiNetCluster realizes a general multi-view learning approach by implementing manifold alignment/warping to combine multiple views into a common latent subspace for further analysis, *i.e*. clustering. Previous studies have emphasized the importance of multiview learning in heterogenous biological data (Li et al., 2016a) or discussed different methods realizing multiview learning (Sun, 2013; Xu et al., 2013) but, to the best of our knowledge, very few of them (Colom-Tatch and Theis, 2018; Li et al., 2016b) regarded manifold alignment as such a method. In our approach, manifold alignment is considered to be a natural and effective method for multiview representation learning. It follows the regularized Empirical Risk Minimization principle (Vapnik, 1992), incorporating in its empirical loss multiple inputs (*e.g*. gene expression profiles), hypothesis spaces, and regularization terms controlling the complexity (or the smoothness) of the solution in the intrinsic geometry of data manifolds (see Equation 1 in Methods).

ManiNetCluster can be used as general purpose to study other biological networks with additional linkage types such as protein-protein interactions. One possible application is the single cell. Increasing single cell data enable identification of interactions among various cell types and seeing how cell types contribute to the phenotypes at the tissue level such as tissue gene expression. Moreover, nonlinearity has been found to widely exist among cell interactions. Thus, ones can also apply this method to single cell gene networks and find out the genomic functional linkages across cell types, providing potential novel insights on cell type interactions.

## Acknowledgments

This work was supported by a Stony Brook University/Brookhaven National Laboratory seed grant to D.W. and I.K.B. I.K.B. was also supported by the Office of Biological and Environmental Research of the Department Of Energy.

## A. Construction of WGCNA and other clustering methods

To compare ManiNetCluster clusterings with WGCNA clustering, we also construct the weighted gene co-expression networks and clustering as follows:

We constructed the gene co-expression networks by connecting all possible gene pairs by edges whose weights are the combination of Pearson correlations and Euclidean distance of their time-series gene expression profiles. The reason for this combination is that Pearson correlations well capture the “shape” of the data (up or down of the expression) while Euclidean distance well capture the “scale” of the data (low or high of the expression). First, we constructed a similarity matrix *S* of a dataset *X* as follows (Hughitt, 2016):

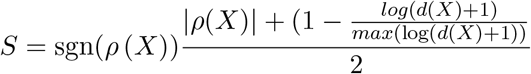

where *ρ*(*X*) depicts the pairwise Pearson correlation and *d*(*X*) depicts the pairwise Euclidean distance of the input dataset. The first term of the equation is the sign of the Pearson correlation, preserving the sign of the interaction. The second terms combine the Pearson correlation and the “Euclidean closeness”, which is the log inverse Euclidean distance. The result *S*, measuring the similarity between two genes, is a number, ranging from −1 to 1, indicating the strength of correlation and its sign of interaction, *i.e*. positive or negative (Hughitt, 2016).

Next, we construct the adjacency matrix from the similarity matrix. We use the power transformation, as suggested by Zhang et al. (Zhang and Horvath, 2005) to reduce the number of spurious correlations in the data and to transform the network into a scale-free topology (Zhang and Horvath, 2005). The resulted adjacency matrix *A* = {*a*_ij_} is computed from similarity matrix *S* = {*s*_ij_} as follows:

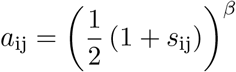

The gene co-expression networks were then clustered into modules by using the *cut-treeDynamicTree* function in WGCNA (weighted correlation network analysis) R package (Langfelder and Horvath, 2008).

Clustering results of *k*-means, hierarchical clustering, and expectation maximization is obtained directly from functions *kmeans(), cutree(hclust())*, and *Mclust()* respectively in R packages *cluster* (Maechler et al., 2012) and *Mclust* (Scrucca et al., 2016). These methods are not for simultaneous clustering, so we perform these on light period genes and dark period genes separately. The number of cluster (*k*) is 34 for light period genes and 30 for dark period genes.

## References

Charu C Aggarwal, Alexander Hinneburg, and Daniel A Keim. On the surprising behavior of distance metrics in high dimensional space. In International conference on database theory, pages 420–434. Springer, 2001.

Ayelet Alpert, Lindsay S. Moore, Tania Dubovik, and Shai S. Shen-Orr. Alignment of single-cell trajectories to compare cellular expression dynamics. Nature methods, 15(4): 267–267, 2018.

M. Belkin and P. Niyogi. Laplacian eigenmaps for dimensionality reduction and data representation. Neural Computation, 15(6):1373–1396, 2003. ISSN 0899-7667. doi: Doi10.1162/089976603321780317. URL <GotoISI>://wos:000182530600005.

Mikhail Belkin, Partha Niyogi, and Vikas Sindhwani. Manifold regularization: A geometric framework for learning from labeled and unlabeled examples. Journal of machine learning research, 7(Nov):2399–2434, 2006.

Scott L Carter, Christian M Brechbhler, Michael Griffin, and Andrew T Bond. Gene co-expression network topology provides a framework for molecular characterization of cellular state. Bioinformatics, 20(14):2242–2250, 2004. ISSN 1460-2059.

Susan E Celniker, Laura AL Dillon, Mark B Gerstein, Kristin C Gunsalus, Steven Henikoff, Gary H Karpen, Manolis Kellis, Eric C Lai, Jason D Lieb, and David M MacAlpine. Unlocking the secrets of the genome. Nature, 459(7249):927, 2009. ISSN 1476-4687.

M Colom-Tatch and FJ Theis. Statistical single cell multi-omics integration. Current Opinion in Systems Biology, 7:54–59, 2018. ISSN 2452-3100.

Ehsan Elhamifar and Ren Vidal. Sparse manifold clustering and embedding. In Advances in neural information processing systems, pages 55–63, 2011.

Charles Fefferman, Sanjoy Mitter, and Hariharan Narayanan. Testing the manifold hypothesis. Journal of the American Mathematical Society, 29(4):983–1049, 2016.

Jihun Ham, Daniel D Lee, and Lawrence K Saul. Semisupervised alignment of manifolds. In AISTATS, pages 120–127, 2005.

M. L. Heinnickel and A. R. Grossman. The greencut: re-evaluation of physiological role of previously studied proteins and potential novel protein functions. Photosynth Res, 116(2-3):427–36, 2013. ISSN 1573-5079 (Electronic) 0166-8595 (Linking). doi: 10.1007/ s11120-013-9882-6. URL https://www.ncbi.nlm.nih.gov/pubmed/23873414.

V Keith Hughitt. Supplemental file: sandfly co-expression cluster analysis. 2016.

Pablo A. Jaskowiak, Ricardo J. G. B. Campello, and Ivan G. Costa. On the selection of appropriate distances for gene expression data clustering. In BMC bioinformatics, volume 15, pages S2–S2, 2014.

Karel Kalecky and Young-Rae Cho. Primalign: Pagerank-inspired markovian alignment for large biological networks. Bioinformatics, 34(13):i537–i546, 2018. ISSN 1367-4803.

S. J. Karpowicz, S. E. Prochnik, A. R. Grossman, and S. S. Merchant. The greencut2 resource, a phylogenomically derived inventory of proteins specific to the plant lineage. J Biol Chem, 286(24):21427–39, 2011. ISSN 1083-351X (Electronic) 0021-9258 (Linking). doi: 10.1074/jbc.M111.233734. URL https://www.ncbi.nlm.nih.gov/pubmed/21515685.

H. Kirst, J. G. Garcia-Cerdan, A. Zurbriggen, and A. Melis. Assembly of the light-harvesting chlorophyll antenna in the green alga chlamydomonas reinhardtii requires expression of the tla2-cpftsy gene. Plant Physiol, 158(2):930–45, 2012. ISSN 1532-2548 (Electronic) 0032-0889 (Linking). doi: 10.1104/pp.111.189910. URL https://www.ncbi.nlm.nih.gov/pubmed/22114096.

Peter Langfelder and Steve Horvath. Wgcna: an r package for weighted correlation network analysis. BMC bioinformatics, 9(1):559–559, 2008.

Neil D Lawrence. A unifying probabilistic perspective for spectral dimensionality reduction: Insights and new models. Journal of Machine Learning Research, 13(May):1609–1638, 2012.

Celine Lefebvre, Presha Rajbhandari, Mariano J Alvarez, Pradeep Bandaru, Wei Keat Lim, Mai Sato, Kai Wang, Pavel Sumazin, Manjunath Kustagi, and Brygida C Bisikirska. A human bcell interactome identifies myb and foxm1 as master regulators of proliferation in germinal centers. Molecular systems biology, 6(1):377, 2010. ISSN 1744-4292.

Yifeng Li, Fang-Xiang Wu, and Alioune Ngom. A review on machine learning principles for multi-view biological data integration. Briefings in bioinformatics, 19(2):325–340, 2016a. ISSN 1467-5463.

Yingming Li, Ming Yang, and Zhongfei Zhang. Multi-view representation learning: A survey from shallow methods to deep methods. arXiv preprint arXiv:1610.01206, 2016b.

Chung-Shou Liao, Kanghao Lu, Michael Baym, Rohit Singh, and Bonnie Berger. Isorankn: spectral methods for global alignment of multiple protein networks. Bioinformatics, 25 (12):i253–i258, 2009. ISSN 1460-2059.

Qi Liao, Changning Liu, Xiongying Yuan, Shuli Kang, Ruoyu Miao, Hui Xiao, Guoguang Zhao, Haitao Luo, Dechao Bu, and Haitao Zhao. Large-scale prediction of long non-coding rna functions in a codingnon-coding gene co-expression network. Nucleic acids research, 39(9):3864–3878, 2011. ISSN 1362-4962.

Martin Maechler, Peter Rousseeuw, Anja Struyf, Mia Hubert, and Kurt Hornik. Cluster: cluster analysis basics and extensions. R package version, 1(2):56, 2012.

John S Mattick, Ryan J Taft, and Geoffrey J Faulkner. A global view of genomic informationmoving beyond the gene and the master regulator. Trends in genetics, 26(1):21–28, 2010. ISSN 0168-9525.

S. S. Merchant, S. E. Prochnik, O. Vallon, E. H. Harris, S. J. Karpowicz, G. B. Witman, A. Terry, A. Salamov, L. K. Fritz-Laylin, L. Marechal-Drouard, W. F. Marshall, L. H. Qu, D. R. Nelson, A. A. Sanderfoot, M. H. Spalding, V. V. Kapitonov, Q. Ren, P. Ferris, E. Lindquist, H. Shapiro, S. M. Lucas, J. Grimwood, J. Schmutz, P. Cardol, H. Cerutti, G. Chanfreau, C. L. Chen, V. Cognat, M. T. Croft, R. Dent, S. Dutcher, E. Fernandez, H. Fukuzawa, D. Gonzalez-Ballester, D. Gonzalez-Halphen, A. Hallmann, M. Hanikenne, M. Hippler, W. Inwood, K. Jabbari, M. Kalanon, R. Kuras, P. A. Lefebvre, S. D. Lemaire, A. V. Lobanov, M. Lohr, A. Manuell, I. Meier, L. Mets, M. Mittag, T. Mittelmeier, J. V. Moroney, J. Moseley, C. Napoli, A. M. Nedelcu, K. Niyogi, S. V. Novoselov, I. T. Paulsen, G. Pazour, S. Purton, J. P. Ral, D. M. Riano-Pachon, W. Riekhof, L. Rymarquis, M. Schroda, D. Stern, J. Umen, R. Willows, N. Wilson, S. L. Zimmer, J. Allmer, J. Balk, K. Bisova, C. J. Chen, M. Elias, K. Gendler, C. Hauser, M. R. Lamb, H. Ledford, J. C. Long, J. Minagawa, M. D. Page, J. Pan, W. Pootakham, S. Roje, A. Rose, E. Stahlberg, A. M. Terauchi, P. Yang, S. Ball, C. Bowler, C. L. Dieckmann, V. N. Gladyshev, P. Green, R. Jorgensen, S. Mayfield, B. Mueller-Roeber, S. Rajamani, R. T. Sayre, P. Brokstein, et al. The chlamydomonas genome reveals the evolution of key animal and plant functions. Science, 318(5848):245–50, 2007. ISSN 1095-9203 (Electronic) 0036-8075 (Linking). doi: 10.1126/science.1143609. URL https://www.ncbi.nlm.nih.gov/pubmed/17932292.

Ha Quang Minh and Vikas Sindhwani. Vector-valued manifold regularization. In ICML, pages 57–64. Citeseer, 2011.

Ha Quang Minh, Loris Bazzani, and Vittorio Murino. A unifying framework in vector-valued reproducing kernel hilbert spaces for manifold regularization and co-regularized multi-view learning. The Journal of Machine Learning Research, 17(1):769–840, 2016. ISSN 1532-4435.

Manikandan Narayanan, Adrian Vetta, Eric E. Schadt, and Jun Zhu. Simultaneous clustering of multiple gene expression and physical interaction datasets. PLoS computational biology, 6(4):e1000742–e1000742, 2010.

G. J. Pazour, N. Agrin, J. Leszyk, and G. B. Witman. Proteomic analysis of a eukaryotic cilium. J Cell Biol, 170(1):103–13, 2005. ISSN 0021-9525 (Print) 0021-9525 (Linking). doi: 10.1083/jcb.200504008. URL https://www.ncbi.nlm.nih.gov/pubmed/15998802.

V. Saraph and T. Milenkovic. Magna: Maximizing accuracy in global network alignment. Bioinformatics, 30(20):2931–40, 2014. ISSN 1367-4811 (Electronic) 1367-4803 (Linking). doi: 10.1093/bioinformatics/btu409. URL https://www.ncbi.nlm.nih.gov/pubmed/25015987.

A. Sawyer and M. Winkler. Evolution of chlamydomonas reinhardtii ferredoxins and their interactions with [fefe]-hydrogenases. Photosynth Res, 134(3):307–316, 2017. ISSN 1573-5079 (Electronic) 0166-8595 (Linking). doi: 10.1007/s11120-017-0409-4. URL https://www.ncbi.nlm.nih.gov/pubmed/28620699.

Luca Scrucca, Michael Fop, T Brendan Murphy, and Adrian E Raftery. mclust 5: Clustering, classification and density estimation using gaussian finite mixture models. The R journal, 8(1):289, 2016.

Vikas Sindhwani and David S Rosenberg. An rkhs for multi-view learning and manifold co-regularization. In Proceedings of the 25th international conference on Machine learning, pages 976–983. ACM, 2008. ISBN 1605582050.

Archana Singh, Avantika Yadav, and Ajay Rana. K-means with three different distance metrics. International Journal of Computer Applications, 67(10), 2013. ISSN 0975-8887.

Rohit Singh, Jinbo Xu, and Bonnie Berger. Global alignment of multiple protein interaction networks with application to functional orthology detection. Proceedings of the National Academy of Sciences, 2008. ISSN 0027-8424.

Shiliang Sun. A survey of multi-view machine learning. Neural Computing and Applications, 23(7-8):2031–2038, 2013. ISSN 0941-0643.

O. Thimm, O. Blasing, Y. Gibon, A. Nagel, S. Meyer, P. Kruger, J. Selbig, L. A. Muller, S. Y. Rhee, and M. Stitt. Mapman: a user-driven tool to display genomics data sets onto diagrams of metabolic pathways and other biological processes. Plant J, 37(6):914–39, 2004. ISSN 0960-7412 (Print) 0960-7412 (Linking). URL https://www.ncbi.nlm.nih.gov/pubmed/14996223.

Vladimir Vapnik. Principles of risk minimization for learning theory. In Advances in neural information processing systems, pages 831–838, 1992.

V. Vijayan, V. Saraph, and T. Milenkovic. Magna++: Maximizing accuracy in global network alignment via both node and edge conservation. Bioinformatics, 31(14):2409–11, 2015. ISSN 1367-4811 (Electronic) 1367-4803 (Linking). doi: 10.1093/bioinformatics/ btv161. URL https://www.ncbi.nlm.nih.gov/pubmed/25792552.

Hoa Trong Vu, Clifton Carey, and Sridhar Mahadevan. Manifold warping: Manifold alignment over time. In AAAI, volume 1, pages 8–8, 2012.

Chang Wang and Sridhar Mahadevan. A general framework for manifold alignment. In AAAI fall symposium: manifold learning and its applications, pages 53–58, 2009.

Weiran Wang, Raman Arora, Karen Livescu, and Jeff Bilmes. On deep multi-view representation learning. In International Conference on Machine Learning, pages 1083–1092, 2015.

Joshua D. Welch, Alexander J. Hartemink, and Jan F. Prins. Matcher: manifold alignment reveals correspondence between single cell transcriptome and epigenome dynamics. Genome biology, 18(1):138–138, 2017.

Chang Xu, Dacheng Tao, and Chao Xu. A survey on multi-view learning. arXiv preprint arXiv:1304.5634, 2013.

Koon-Kiu Yan, Daifeng Wang, Joel Rozowsky, Henry Zheng, Chao Cheng, and Mark Gerstein. Orthoclust: an orthology-based network framework for clustering data across multiple species. Genome biology, 15(8):R100, 2014. ISSN 1474-760X.

Koon-Kiu Yan, Daifeng Wang, Anurag Sethi, Paul Muir, Robert Kitchen, Chao Cheng, and Mark Gerstein. Cross-disciplinary network comparison: matchmaking between hairballs. Cell systems, 2(3):147–157, 2016. ISSN 2405-4712.

Bo Yang, Xiao Fu, Nicholas D. Sidiropoulos, and Mingyi Hong. Towards k-means-friendly spaces: Simultaneous deep learning and clustering. arXiv preprint arXiv:1610.04794, 2016.

Yang Yang, Leng Han, Yuan Yuan, Jun Li, Nainan Hei, and Han Liang. Gene co-expression network analysis reveals common system-level properties of prognostic genes across cancer types. Nature communications, 5:3231, 2014. ISSN 2041-1723.

Andy M Yip and Steve Horvath. Gene network interconnectedness and the generalized topological overlap measure. BMC bioinformatics, 8(1):22, 2007. ISSN 1471-2105.

Xinghuo Zeng, Matthew J. Nesbitt, Jian Pei, Ke Wang, Ismael A. Vergara, and Nansheng Chen. Orthocluster: a new tool for mining synteny blocks and applications in comparative genomics. In Proceedings of the 11th international conference on Extending database technology: Advances in database technology, pages 656–667, 2008.

Bin Zhang and Steve Horvath. A general framework for weighted gene co-expression network analysis. Statistical applications in genetics and molecular biology, 4(1), 2005.

James Matt Zones, Ian K. Blaby, Sabeeha S. Merchant, and James G. Umen. High-resolution profiling of a synchronized diurnal transcriptome from chlamydomonas reinhardtii reveals continuous cell and metabolic differentiation. The Plant Cell, pages tpc–15, 2015.

